# Optimizing Sample Preparation for Direct Nanopore Sequencing to Enable Rapid Pathogen and Antimicrobial Resistance Profiling in Bovine Mastitis

**DOI:** 10.1101/2025.08.19.671071

**Authors:** Crystal Chapagain, Abdolrahman Khezri, Jawad Ali, Marit Smistad, Liv Synnøve Sølverød, Rafi Ahmad

## Abstract

Long-read metagenomic sequencing allows for the rapid, culture-independent, and accurate identification of causative pathogens and antimicrobial resistance (AMR) profiles, supporting precise antibiotic use and reducing the spread of resistance. However, its application to mastitis milk is challenging due to the complex milk matrix, low bacterial count, and high somatic cell content. This study primarily aimed to further optimize our previously developed direct sequencing protocol for milk samples from mastitis cases. Additional optimizations included combining centrifugation, gradient centrifugation, and fat fraction treatment with Tween 20 and citric acid. Subsequently, four DNA extraction kits (Blood and Tissue, Molysis Complete5, HostZero, and SPINeasy Host depletion) were evaluated for their ability to remove host DNA and enrich bacterial DNA for long-read sequencing with Oxford Nanopore technologies. qPCR was used to quantify bacterial and bovine DNA, allowing comparison of host depletion efficiency among the kits.

Our results show that simple centrifugation effectively concentrates bacterial cells, removing the need for chemical treatments. The HostZero kit consistently produced higher DNA yields, better DNA integrity, and more effective host DNA depletion. Using nanopore sequencing, both Gram-positive and Gram-negative mastitis pathogens, along with their AMR genes, were successfully detected. Overall, this study underscores the importance of an effective DNA extraction method for the direct sequencing of mastitis milk samples. Additionally, our findings support the potential of direct metagenomic sequencing as a rapid, culture-free approach for identifying mastitis pathogens and their resistance profiles.

## Introduction

Mastitis is an inflammation of the mammary gland usually caused by various pathogens, invading the udder tissue, and occasionally by mechanical or chemical trauma (Malcata et al., 2020; Ramuada et al., 2024). It is a costly infectious disease, estimated to cost €16–26 billion annually to the global dairy industry (University of Glasgow, 2016) due to reduced milk production, milk wastage, treatment costs, early culling, and, in severe cases, mortality (Abebe et al., 2016). In Norway, records from the Norwegian Dairy Herd Recording System (NDHRS) indicate that mastitis accounts for more than one-third of all reported diseases in dairy cows and is the leading cause of antibiotic use (TINE, 2025).

Although over 134 pathogens, including bacteria, viruses, mycoplasma, yeasts, and algae, have been linked to bovine mastitis, bacteria cause about 95% of all cases (Zigo et al., 2021). *Staphylococcus aureus* is the leading cause of both clinical and subclinical mastitis in Norway. Other commonly identified pathogens include non-*aureus Staphylococci and Mammaliicocci* (NASM), *Escherichia coli*, and *Streptococcus* species. (Smistad et al., 2023). Mastitis is typically categorized into clinical and subclinical types. Clinical mastitis is further divided into mild, moderate, and severe levels depending on symptom intensity. It is marked by visible changes in the milk, such as clots, discoloration, blood, or a watery look, along with signs of inflammation in the udder, like swelling, heat, redness, and pain. Conversely, subclinical mastitis often shows no obvious symptoms but can be identified through increased somatic cell count (SCC) and bacterial cultures. It acts as a reservoir for pathogen spread within the herd (Urrutia-Angulo et al., 2024).

Mastitis is primarily treated with antimicrobials based on the clinical diagnosis report. The current gold standard for diagnosis involves culturing milk samples, identifying pathogens through biochemical tests or MALDI-TOF mass spectrometry, and performing culture-based antibiotic susceptibility testing. This traditional method takes 3 to 5 days and has notable limitations in mastitis diagnosis because the specificity of culture-based detection is relatively low (Jung et al., 2019; Chamchoy et al., 2022; Smistad et al., 2022), either due to the presence of mixed bacterial populations or the absence of detectable growth. The delayed information on the causative agent and the correct antibiotic often leads to the empirical use of broad-spectrum antibiotics (Malcata et al., 2020), contributing to the emergence of AMR, a growing global public health concern (WHO, 2016). Overusing antimicrobials for an extended period can lead to drug residues in milk, contributing to the spread of resistance to humans and resulting in additional economic losses. Furthermore, inappropriate or delayed treatment compromises animal welfare by causing prolonged pain, discomfort, and reduced quality of life (Ruegg, 2017). Therefore, developing rapid and reliable methods to identify mastitis-causing pathogens and their resistance profiles is essential for improving diagnostic accuracy and promoting the responsible use of antibiotics.

Molecular techniques based on PCR offer high sensitivity, but they can only detect a limited number of pathogens and antimicrobial-resistant genes (ARGs) (Yamin et al., 2023). In recent years, metagenomics and long-read sequencing technologies have become faster, more accurate, and affordable, gaining significant attention as powerful tools for more comprehensive and unbiased diagnostic solutions (Satam et al., 2023). Many studies have used Oxford Nanopore sequencing technology to identify pathogens and ARGs in human clinical samples, such as blood (Ali et al., 2024; Gu et al., 2025), urine (Liu et al., 2023b; Bellankimath et al., 2024), bronchoalveolar lavage fluid (Li et al., 2025), as well as in bovine milk samples (Ahmadi et al., 2023; Usui et al., 2023). A significant challenge in sequencing-based diagnostics is the abundance of host DNA, which can shadow the microbial genome in the milk samples. This is particularly true for mastitis milk due to its high fat and somatic cell content, which easily exceeds 200,000 cells/ml, even in subclinical mastitis (Liu et al., 2023a).

In our previous work, we described a culture- and amplification-independent sequencing approach to identify pathogens and antibiotic resistance genes in mastitis milk samples, which could potentially reduce diagnostic time to 5–9 hours. This study aimed to optimize the sample treatment protocols before DNA extraction to remove the matrix without affecting the viability of bacterial cells from clinical mastitis milk samples. Additionally, we compared four commercial DNA extraction kits for effective host depletion and microbial DNA isolation from mastitis milk samples, suitable for Oxford Nanopore metagenomic sequencing to identify the causative pathogens and their AMR profiles.

## METHODOLOGY

### Milk samples

In this study, 10 quarter milk samples from 10 different cows diagnosed with clinical mastitis caused by various gram-positive (*S. aureus, Streptococcus dysgalactiae, Streptococcus uberis*) and gram-negative (*E. coli*) bacteria were provided by TINE SA from the routine mastitis diagnostics. Initial bacterial identification was performed at TINE’s Mastitis Laboratory (Molde, Norway), using overnight culturing followed by MALDI TOF mass spectrometry. To minimize the freezing effect on genomic material, 30% glycerol (v/v) was added to the samples. The frozen samples were shipped to the INN laboratory in Hamar, Norway, and kept at -20 °C. On the day of the experiment, samples were thawed at room temperature and re-cultured in Brain Heart Infusion (BHI) agar (15 g/L Agar, 37 g/L BHI Broth, VWR Life Science, USA) to determine the CFU/mL values post-freezing. The experimental design overview is presented in Figure 1.

**Figure 1.**
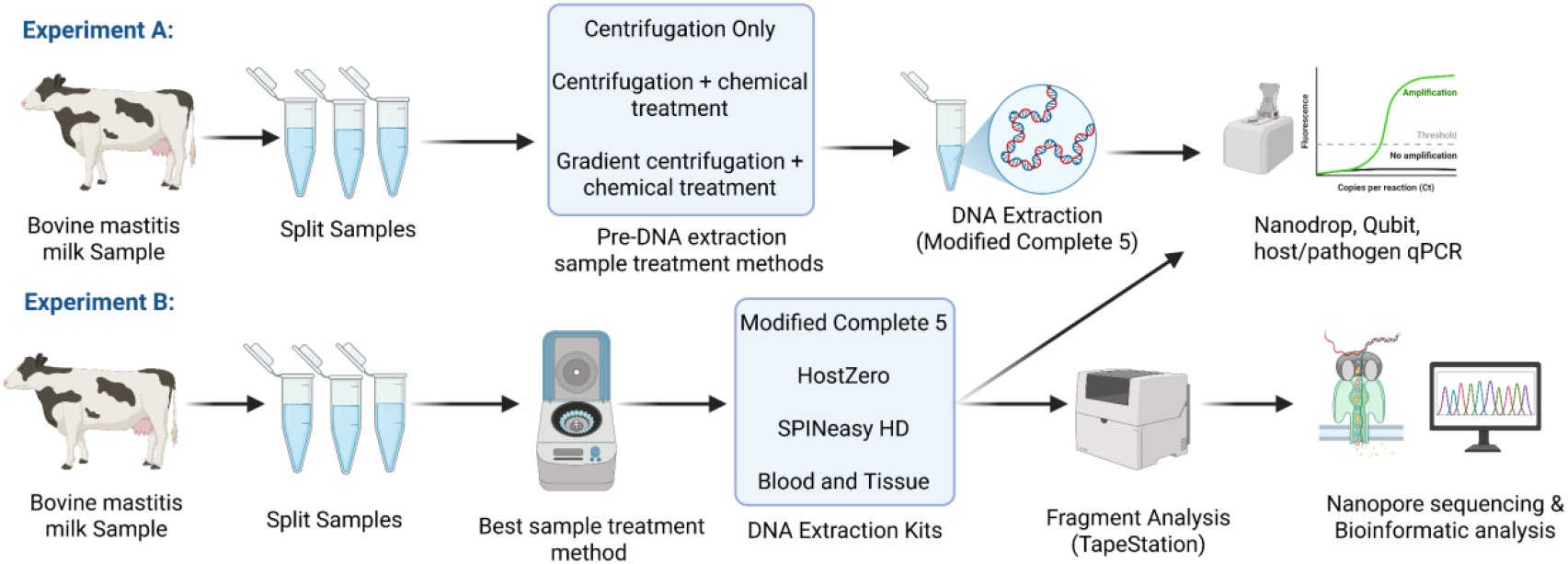
A graphical overview of the experimental design. *Experiment A*: Evaluation of different pre-DNA extraction sample treatment methods; *Experiment B*: The optimized sample treatment method from Experiment A was employed for DNA extraction using four different commercial kits, followed by qPCR, TapeStation analysis, nanopore sequencing, and bioinformatics analysis.

### Experimental design

#### Experiment A: Optimizing the pre-DNA extraction sample treatment

In our previous study (Ahmadi et al., 2023), the MolYsis™ Complete5 kit (Molzym, Bremen, Germany) demonstrated optimal performance when an additional centrifugation step was included before proceeding with the manufacturer’s protocol for DNA isolation. In this study, three different methods were analyzed to optimize the recovery of bacterial cells that may become trapped in fat globules and are often lost in the supernatant during centrifugation. The workflow of these three methods is presented in (Supplementary Figure 1). The pre-DNA extraction sample treatment optimization was performed using three representative milk samples with varying bacterial loads: high (∼10^7^ CFU/mL), medium (∼10^5^ CFU/mL), and low (∼ 10^3^ CFU/mL). Each milk sample was divided into three 1 mL aliquots and subjected to the following pretreatment methods.

##### Method 1 – Centrifugation only

Milk samples were centrifuged at 4500 *x g* for 20 minutes at 4°C to separate the fat and whey layers from the cellular components. The upper fat and whey fractions were carefully removed, and the remaining pellet was retained. To reduce residual components, the pellet was washed with 1 mL phosphate-buffered saline (PBS) and centrifuged at 13000 *x g* for 1 minute (Ahmadi et al., 2023). The washing step was performed twice.

##### Method 2 -Centrifugation followed by chemical treatment

Initial centrifugation was performed as described in Method 1 (4,500 × g for 20 minutes at 4 °C). The resulting pellet was kept on ice, while the supernatant, comprising the fat and whey layers, was subjected to further processing. To disrupt protein and fat components and release any bacterial cells potentially trapped within them, the supernatant was incubated with 0.1% Tween 20 and 2% citric acid at room temperature for 15 minutes (Duarte and Porcellato, 2024). The treated layer was centrifuged at 8000 *x g* for 10 minutes at 4 ^0^C, and the pellet was combined with the original pellet from the initial centrifugation. The combined pellet was subsequently washed twice with PBS as described in Method 1.

##### Method 3 -Gradient centrifugation combined with chemical Treatment

An equal volume of Percoll solution (1.050 g/ml) was added to the milk sample to create a density gradient, followed by centrifugation at 4500 *x g* for 15 minutes at room temperature (Meisel et al., 2011). The supernatant was carefully transferred to a clean microcentrifuge tube, then chemically treated and combined with the initial pellet. The combined pellet was washed twice and resuspended in PBS.

Following sample treatment, a 100 µL aliquot of the final supernatant from all three tested conditions was plated on BHI agar (VWR Life Sciences, USA) and incubated at 37 ^0^C for 24 hours. The CFUs were counted to assess potential bacterial loss compared to the initial CFUs of the selected samples. The resulting pellets, presumed to contain concentrated bacterial cells, were suspended in 1 mL PBS for DNA extraction. The modified protocol (Mol Com5_cent-nuc_) of the MolYsis™ Complete5 kit, containing an additional micrococcal nuclease treatment after host depletion, was used for extracting DNA as previously described (Ahmadi et al., 2023). The extracted DNA was assessed using qPCR, with primers specific to *S. aureus* and bovine (Supplementary Table 1). The reaction condition and thermal profile for qPCR were as described below.

**Table 1.** An overview of DNA quality and sequencing metrics for mastitis milk samples using two DNA isolation kits. The DIN value for the *S. aureus* Mol Com5 sample was not generated by TapeStation, indicating very poor DNA integrity in the sample. Mol Com 5 and HostZero are short forms for Mol Com5 and HostZero, respectively. For *S. aureus* Mol Com5, two values for DNA fragment size represent the two peaks identified by the TapeStation analysis, with the accompanying percentages indicating the proportion of total DNA corresponding to each fragment size.

| Ground truth pathogen in the mastitis milk sample | DNA Isolation Kit | Qubit 4 | TapeStation |  | NanoStat |  |  |  | Blastn and Minimap2 |  |  |
| --- | --- | --- | --- | --- | --- | --- | --- | --- | --- | --- | --- |
|  |  | DNA yield (ng) | DNA fragment size ( | DIN value | Total Number of passed reads | Read N50 | Mean read quality (Q-score) | Mean read length | Reads mapped with targeted pathogens (%) | Reads mapped with Un-targeted pathogens (%) | Host Reads (bovine) (%) |
| *S. aureus* | Mol Com5 | 68.4 | 2140(53.11%), 3829 (36.45%) | - | 9550 | 4631 | 11.4 | 4095 | 77.2 | 0.08 | 22.72 |
|  | HostZero | 46.75 | 59474 | 7.9 | 115404 | 3969 | 11.6 | 3594 | 87.96 | 0.1 | 11.93 |
| *E. coli* | Mol Com5 | 89.2 | 16555 | 1.8 | 14005 | 3778 | 10.4 | 3102 | 75.75 | 9.45 (5% *S. aureus* reads) | 14.78 |
|  | HostZero | 350 | 23655 | 7.4 | 101754 | 3707 | 10.6 | 3324 | 82.09 | 5.13 | 12.77 |

#### Experiment B: Evaluation of DNA extraction kits

##### Direct DNA extraction from milk samples

Four commercial DNA extraction kits were evaluated for their effectiveness in isolating microbial DNA from mastitis-infected milk, including three kits specifically designed for the selective depletion of host DNA. The kits tested were: a modified version of the MolYsis™ Complete 5 kit (Mol Com5_cent-nuc_) as described by Ahmadi et al. (2023), HostZERO Microbial DNA Kit (Zymo Research), SPINeasy**®** Host Depletion Microbial DNA Kit (MP Biomedicals), and DNeasy Blood & Tissue Kit (Qiagen), hereafter Mol Com5, HostZero, SPINeasy and Blood and Tissue, respectively.

The study utilized five clinical mastitis milk samples, each infected with commonly encountered bovine mastitis pathogens, including *S. aureus, S. dysgalactiae*, and *S. uberis*— and *E. coli*. Sample pretreatment was performed according to the previously described “Method 1”. After washing, the pellets were resuspended in sterile PBS, and volumes were adjusted to meet the input requirements specified for each kit. DNA extraction was performed according to the respective manufacturers’ protocols. Final DNA elution was performed using 100 µL of elution buffer for Mol Com5_cent-nuc_ and Blood and Tissue kits, and 50 µL for HostZERO and SPINeasy kits.

### DNA quality assessment

All samples were evaluated for quantity, purity, and fragment length after DNA extraction. DNA concentration was measured using the Qubit High Sensitivity Assay kit and the Qubit 4.0 fluorometer (Invitrogen, USA), following the manufacturer’s protocol. Sample purity was assessed using a Nanodrop ND-1000 spectrophotometer (NanoDrop Technologies, Rockland, DE, United States), which measured the absorption ratios at 260/280 nm and 260/230 nm. The fragment length and integrity of the extracted DNA (one representative gram-positive: *S. aureus* and one gram-negative: *E. coli*) were analyzed using the Agilent 4150 TapeStation System using Genomic DNA ScreenTape Analysis (Agilent Technologies, USA) for the kits that include host depletion mechanisms (Mol Com5, HostZero, and SPINeasy kits).

### qPCR

To determine the relative proportions of bacterial and bovine DNA, qPCR was performed using species-specific primers (Supplementary table 1). Each reaction was conducted in a total volume of 15 µL, containing 3 µL of 5X Hotfire Pol EvaGreen qPCR supermix (Solis BioDyne, Estonia), 0.3 µL each of 10 µM forward and reverse primers, and 1 µL of template DNA. Nucleic acid- and nuclease-free water was used as a substitute for template DNA in negative control reactions.

qPCR amplification was carried out using a 7500 Fast Real-Time PCR system (Invitrogen ™, USA) under the following thermal cycling conditions: initial denaturation at 95 °C for 12 min, followed by 40 cycles of 95 °C for 25 seconds, 60 °C for 45 seconds (data collection stage), and 72 °C for 1 minute.

### MinION library preparation and sequencing

DNA samples from one mastitis milk sample infected with *S. aureus* (gram-positive) infection and one from *E. coli* (gram-negative) infection were selected for sequencing. For each species, DNA was extracted using Mol Com5 and HostZero kits. Before library preparation, DNA samples were purified and concentrated using AMPure XP beads (Beckman Coulter™, USA) to improve purity and yield. Library preparation was performed using the Oxford Nanopore Technologies Rapid PCR Barcoding kit 24 V14 (SQK-RPB114.24), according to the manufacturer’s instructions. Sequencing was performed for over 24 hours using the R10.4.1 flow cell (FLO-MIN 114) mounted on a MinION MK1D device (Oxford Nanopore Technologies).

### Bioinformatic analysis

Raw sequencing reads were generated and base-called in real-time using the ONT MinKNOW GUI software (version 6.0.11) in FAST base-calling mode, with the Dorado basecaller (version 7.4.13). Sequencing read statistics, including read length, read quality, and N50, were assessed using NanoStat v1.4.0 (De Coster et al., 2018).

To identify potential pathogens, reads were mapped using BLASTn against the NCBI Prokaryotic Reference Genomes collection (RefProk). The BLASTn search was performed with the following parameters: word size-28, maximum target sequences-150, and e-value cutoff-0.000001. The following cutoff values were used for bacterial identification: minimum percent identity: 80%, minimum read coverage in alignment: 65%, minimum read length: 200 nt. Sequencing reads that did not meet the specified criteria or failed to align with the RefProk database were classified as non-aligned, primarily representing the host (bovine) derived sequences.

To create an assembly, the reads were first mapped to the reference genome of the top-identified pathogen using Minimap2 v2.29 (Li, 2018). Correctly mapped reads were extracted using SAMtools v1.13 (Danecek et al., 2021) and subsequently used for *de novo* assembly with Flye v2.9.6 (Kolmogorov et al., 2019). The quality and completeness of the assemblies were evaluated using Quast v5.3.0 (Gurevich et al., 2013) and Benchmarking Universal Single-Copy Orthologs (BUSCO) 5.2.2 (Simão et al., 2015). To determine how well the assembly represents the original sequencing data, reads were mapped back to the assembly using Minimap2 v2.29, enabling an assessment of sequencing depth and the contribution of reads to the assembly.

Assembled genomes were used to identify ARGs and virulence factor (VF) genes using ABricate v1.0.1(Seemann, 2025). To identify the ARGs, the NCBI resistance database (Feldgarden et al., 2019), as well as the CARD 2023 database (Alcock et al., 2023), and to identify VF genes, the core VF database (Liu et al., 2019) within ABricate were used.

## RESULTS

### The centrifugation-only method is adequate, and additional fat fraction treatment did not improve bacterial DNA recovery in milk samples

#### Method 1: Centrifugation only

To concentrate the bacterial cells and remove the unwanted milk components before DNA extraction, milk samples were centrifuged at 4500 *x* g for 20 minutes at 4 ^0^C. The CFU count in the supernatant fat and whey layer, which were discarded after centrifugation, indicated an approximate loss of 5% to 18% of bacterial cells (Figure 2A). This loss was inversely related to the initial bacterial load, with greater losses observed in samples with lower bacterial concentrations.

**Figure 2.**
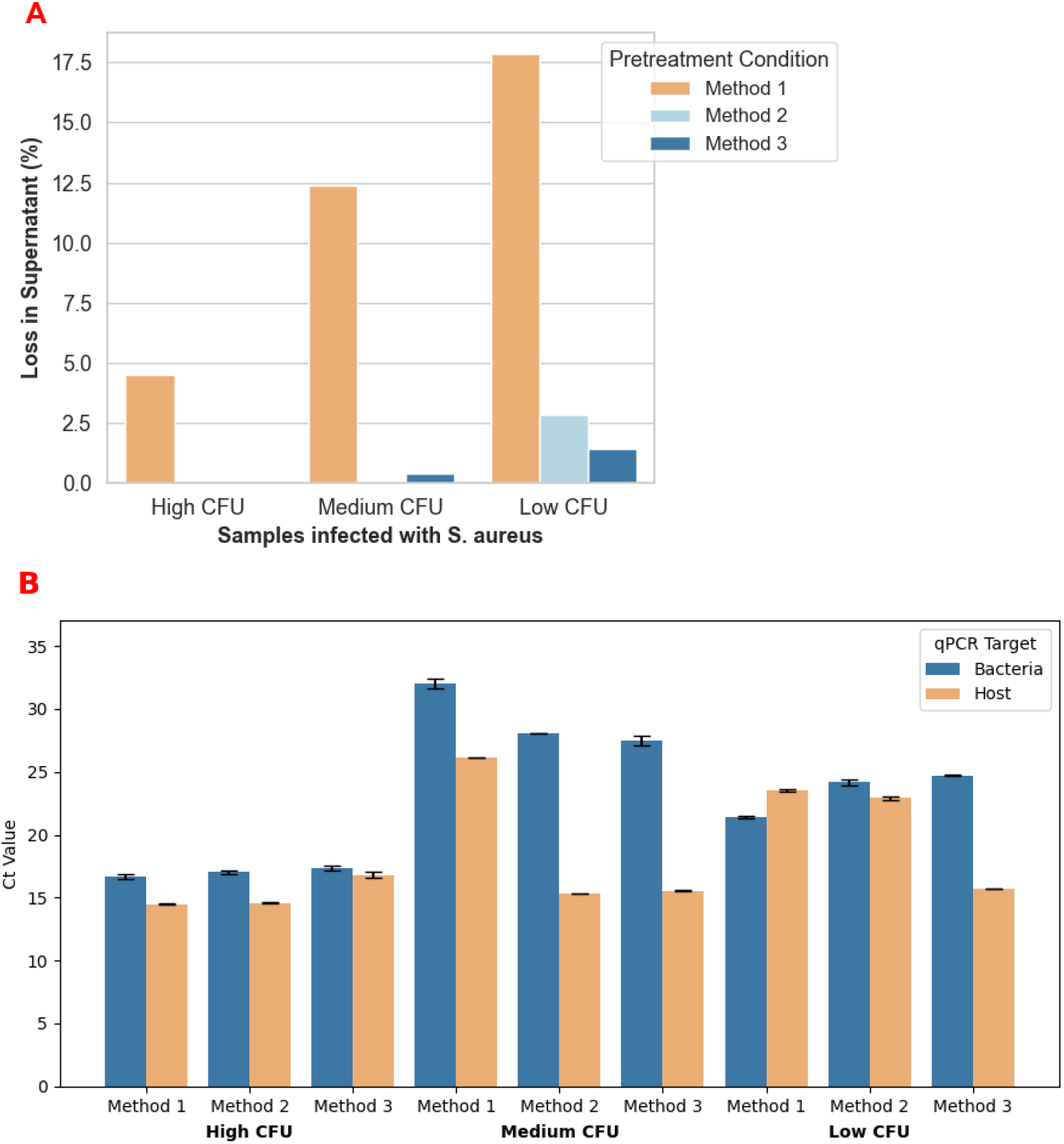
A: Percentage of bacterial loss in the supernatant after three pre-DNA extraction sample eatment methods applied to mastitis milk samples with high, medium, and low bacterial loads. Missing bars indicate values near 0%. B: Ct values following qPCR targeting the nuc gene (Staphylococcus aureus) and BGb gene (bovine) in DNA isolated from mastitis milk samples treated with three pretreatment methods. All qPCR reactions were performed in triplicate, and error bars represent the tandard deviation.

#### Method 2: Centrifugation combined with chemical treatment

To recover bacterial cells trapped in the fat layer after centrifugation (method 1), the fat and whey fractions were additionally treated with 0.1% Tween 20 and 2% citric acid (Supplementary Figure 1), aiming to emulsify the fat and release the associated bacterial cells. Following this chemical treatment and subsequent centrifugation, minimal loss of viable bacteria was observed in the resulting supernatant for samples with high and medium bacterial loads. However, approximately 2.5% of bacterial cell loss was seen in samples with a low initial bacterial concentration.

#### Method 3: Gradient centrifugation combined with chemical treatment

A Percoll gradient was used to improve the separation of bacterial cells trapped in the fat layer (Figure 1). The resulting CFU counts in the supernatant after treatment were negligible across all samples, regardless of their bacterial loads (Figure 2A). These results suggest that bacterial recovery from milk samples is more efficient when chemical treatment is combined with gradient centrifugation.

#### qPCR results

qPCR was performed on the total DNA extracted from treated milk samples to evaluate the effectiveness of different pre-DNA extraction sample treatment strategies in enriching bacterial DNA. The Ct values for both bacterial (*nuc*) and host (*BGb*) targets are presented in Figure 2B. In samples with a high bacterial load, Ct values for the bacterial gene target were comparable across methods, ranging from 16.7 to 17.4. The centrifugation-only method produced the lowest Ct value (16.7), indicating slightly more efficient bacterial DNA recovery. In samples with a low bacterial load, the centrifugation-only method also resulted in a lower bacterial Ct value (21.4) compared to methods 2 (24.2) and 3 (24.7), suggesting better bacterial DNA enrichment. For the medium bacterial load sample, method 1 exhibited the slightly highest bacterial Ct value, indicating lower overall bacterial DNA recovery compared to methods 2 and 3 (Figure 2B). However, the ΔCt (bacterial Ct – host Ct) for method 1 was the lowest (6), compared to 12.8 and 12 for methods 2 and 3, respectively, indicating a more favorable bacterial-to-host DNA ratio, which is critical for downstream bacterial metagenomic analysis.

### HostZero provides superior DNA yield and integrity compared to other host depletion kits

DNA concentration, total yield, and purity ratios (260/280 and 260/230) were evaluated across four commercial DNA extraction kits. The Blood and Tissue kit produced the highest DNA concentration and overall yield in all samples. The HostZero kit provided the most consistent yields across samples infected with different pathogens, with DNA amounts ranging from 13.3 to 350 ng. In contrast, the SPINeasy kit performed the poorest, with DNA concentrations below the detection limit of the Qubit High Sensitivity Assay kit (<0.005 ng/µL) in three samples and very low yields in the remaining two (0.51 ng and 19.7 ng). The Mol Com5 kit also generated relatively low DNA quantities, ranging from 1.22 to 89.2 ng.

Purity ratios varied across different extraction methods and samples. The Blood and tissue kit typically delivered the highest purity, with 260/280 ratios around 1.8 and 260/230 ratios close to the optimal range of 2.0-2.2. In contrast, both the HostZero and Mol Com5 kits consistently exhibited lower 260/280 and 260/230 ratios, indicating the presence of residual protein and salt contamination. The SPINeasy kit often yielded negative or suboptimal 260/230 ratios (see Supplementary Table 2), suggesting potential carryover of solvents or salts.

**Table 2.** Genome assembly and mapping statistics for mastitis milk samples processed with two DNA isolation kits, Mol Com5 and HostZero. Assembly quality was assessed using QUAST and BUSCO. Mapping statistics were obtained using Minimap2 (criteria reported for *Minimap2* were calculated following mapping the reads back to the assembly).

| Ground truth pathogen in the mastitis milk sample | Kit | Quast |  |  |  |  | BUSCO |  |  | Minimap2 |  |  |
| --- | --- | --- | --- | --- | --- | --- | --- | --- | --- | --- | --- | --- |
|  |  | Number of contigs | N50 | AuN | GC% | Genome Fraction | BUSCO Complete % | BUSCO Fragmented % | BUSCO Missing % | Breadth of Cov % | Mean of Cov depth (x) | Unique mapping % |
| *S. aureus* | Mol Com5 | 23 | 285218 | 384734.2 | 32.85 | 90.96 | 51.6 | 30.6 | 17.8 | 99.98 | 11.7 | 99.82 |
|  | HostZero | 4 | 1518621 | 1350486.9 | 32.85 | 92.137 | 98.4 | 1.6 | 0 | 100 | 136.6 | 100 |
| *E. coli* | Mol Com5 | 98 | 40436 | 51880.6 | 51.31 | 68.01 | 43.5 | 40.3 | 16.2 | 99.97 | 8.1 | 83.15 |
|  | HostZero | 23 | 415684 | 457120.8 | 50.84 | 92.312 | 87.9 | 9.7 | 2.4 | 99.98 | 63.2 | 99.96 |

DNA fragmentation analysis using the Agilent TapeStation system revealed that the HostZero kit yielded higher-integrity DNA (DIN 7.4-7.9) compared to Mol Comp5 (0-1.8). However, DNA extracted with the SPINeasy kit had very low concentration and was not within the detectable range of the Genomic DNA Screentape used for the TapeStation; therefore, the fragment size or DNA integrity number could not be determined for this kit (Supplementary Figure 2).

### Host depletion varies across different samples

Among the four kits tested, the Blood and Tissue kit lacks any host depletion mechanism. Meanwhile, HostZero, Mol Com5, and SPINeasy kits are specifically designed for the selective removal of host DNA. Each kit was evaluated across biological replicates, representing milk samples with different pathogens and varying bacterial loads. qPCR was used to quantify bacterial and bovine DNA, and Ct values for each sample are shown in Figure 3.

**Figure 3:**
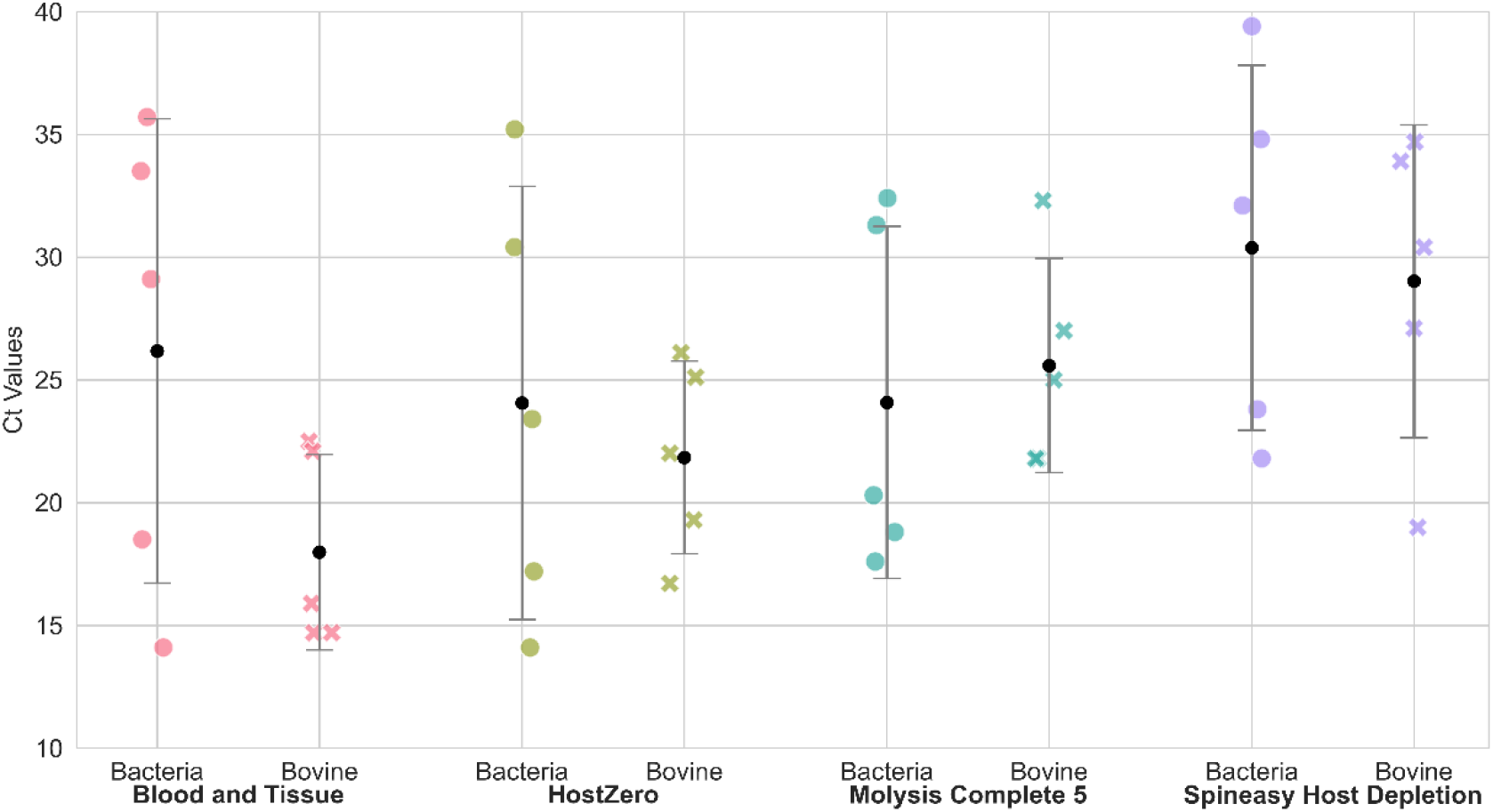
Comparison of Ct values for bacterial and bovine DNA extracted using four different DNA extraction and host depletion kits. Each dot represents the Ct value for one of five samples. Black circles indicate the mean Ct values, with error bars displaying standard deviation. Lower Ct values indicate higher DNA abundance. Higher Ct values for bovine DNA suggest more effective host depletion.

Ct values following the use of the Blood and Tissue kit for bacterial targets ranged from 15 to 35, with consistently low host Ct values, indicating a significant amount of host DNA (Figure 3). Since it lacks a host depletion mechanism, these results are expected to serve as a baseline for comparing the performance of host-depleting kits. The HostZero kit exhibited minimal variation in host Ct values across samples, indicating consistent depletion of host DNA, regardless of bacterial load and pathogen. However, the average host Ct remained lower than the bacterial Ct, indicating that host DNA was still present in abundance. The Mol Com5 kit provided better ΔCt (bacterial Ct – host Ct), which ranged from -6.2 to 5.4, while it ranged from -5.2 to 9.1 for HostZero and -8.6 to 13.1 for the SPINeasy kit (Supplementary table 2). Mol Com5 was the only kit where the average host Ct exceeded the bacterial Ct, indicating effective depletion of host DNA. This kit also produced low bacterial Ct values, supporting efficient bacterial DNA recovery. Although some variation in Ct values was observed across samples, this is expected due to natural biological differences in bacterial load. Overall, Molysis Complete5 showed the strongest performance in selectively enriching bacterial DNA while reducing host background. The DNA extracted with the SPINeasy kit had the highest Ct values for both bacterial and host targets. Higher Ct values for the host target suggest better host depletion; however, the higher Ct values for bacterial targets also indicate that the kit’s DNA isolation capability is compromised.

### Nanopore sequencing & bioinformatic analysis

Based on the evaluation of DNA yield, integrity, and the bacterial-to-host DNA ratio, we selected samples extracted using the Mol Com5 and HostZero kits for downstream nanopore sequencing. The chosen samples include one representative gram-positive and one gram-negative sample. These samples were infected with a high bacterial load (10^7^ CFU/ml) and showed similar DNA yields and Ct values for both bacterial and host DNA across extractions with both kits. Sequencing read length, quality, and taxonomic classification results are summarized in Table 1.

#### Read length, quality, and taxonomy classification

The average read length across all sequenced samples from both kits ranged from approximately 3,100 to 4,100 base pairs (Table 1). Although the *S. aureus* sample processed with the Mol Com5 kit exhibited the highest mean read length among the sequenced samples, it also showed the most significant degree of fragmentation, with notably smaller DNA fragments, as indicated by TapeStation.

Mean read quality (Q-scores) was slightly higher for the *S. aureus* (11.4 to 11.6) sample compared to *E. coli* (10.4 to 10.6) in both extraction methods. The number of reads generated from samples extracted using the HostZero kit was nearly 10 times greater than those from the Mol Com5 kit. After taxonomic classification, 88% and 82% of reads from samples extracted with the HostZero kit were assigned to the target pathogen, compared to 77% and 76% of reads from samples extracted with the Mol Com5 kit. Similarly, about 12% of the reads from the HostZero method were from the host (bovine reads), which was lower than the 23% and 15% of reads classified as host in the samples extracted with Mol Com5 (Table 1).

##### Genome assembly

Both the *S. aureus* and *E. coli* assemblies, generated using the HostZero kit, were of the highest quality among the evaluated methods (Table 2). Assembly quality was assessed using fragmentation metrics, where the HostZero kit produced assemblies with higher N50 values (415684 bp for *E. coli* and 1518621 bp for *S. aureus*) and higher AuN values (457120.8 bp and 1350486.9 bp, respectively), along with a lower number of contigs (23 and 4 contigs for *E. coli* and *S. aureus*), compared to Mol Com5. The superior quality of the assemblies generated with the HostZero kit was further supported by BUSCO analysis, which revealed a higher proportion of complete BUSCO genes (88% for *E. coli* and 98% for *S. aureus*) compared to (44% and 52%, respectively) completeness in the assemblies generated by Mol Com5. In addition, the mean depth of coverage for reads contributing to the final assemblies was remarkably higher with the HostZero kit (63.2 x for *E. coli* and 136.6x for *S. aureus*) than with the Mol Com5 method (8.1x and 11.7x, respectively).

#### AMR gene and VF detection

Using DNA extracted with the HostZero kit, 8 AMR genes and 59 VF in *S. aureus*, as well as all 47 AMR genes and 44 VF in *E. coli*, were detected. In contrast, samples processed with the Mol Com5 kit showed reduced gene recovery: 7 AMR genes were detected in *S. aureus*, and 32 AMR genes were detected in *E. coli*. Similarly, 59 VF genes were detected in *S. aureus* with Mol Com5, and only 37 VF genes were identified in *E. coli* (Figure 4). These results highlight the superior performance of the HostZero kit in recovering both AMR and virulence genes from metagenomic milk samples.

**Figure 4.**
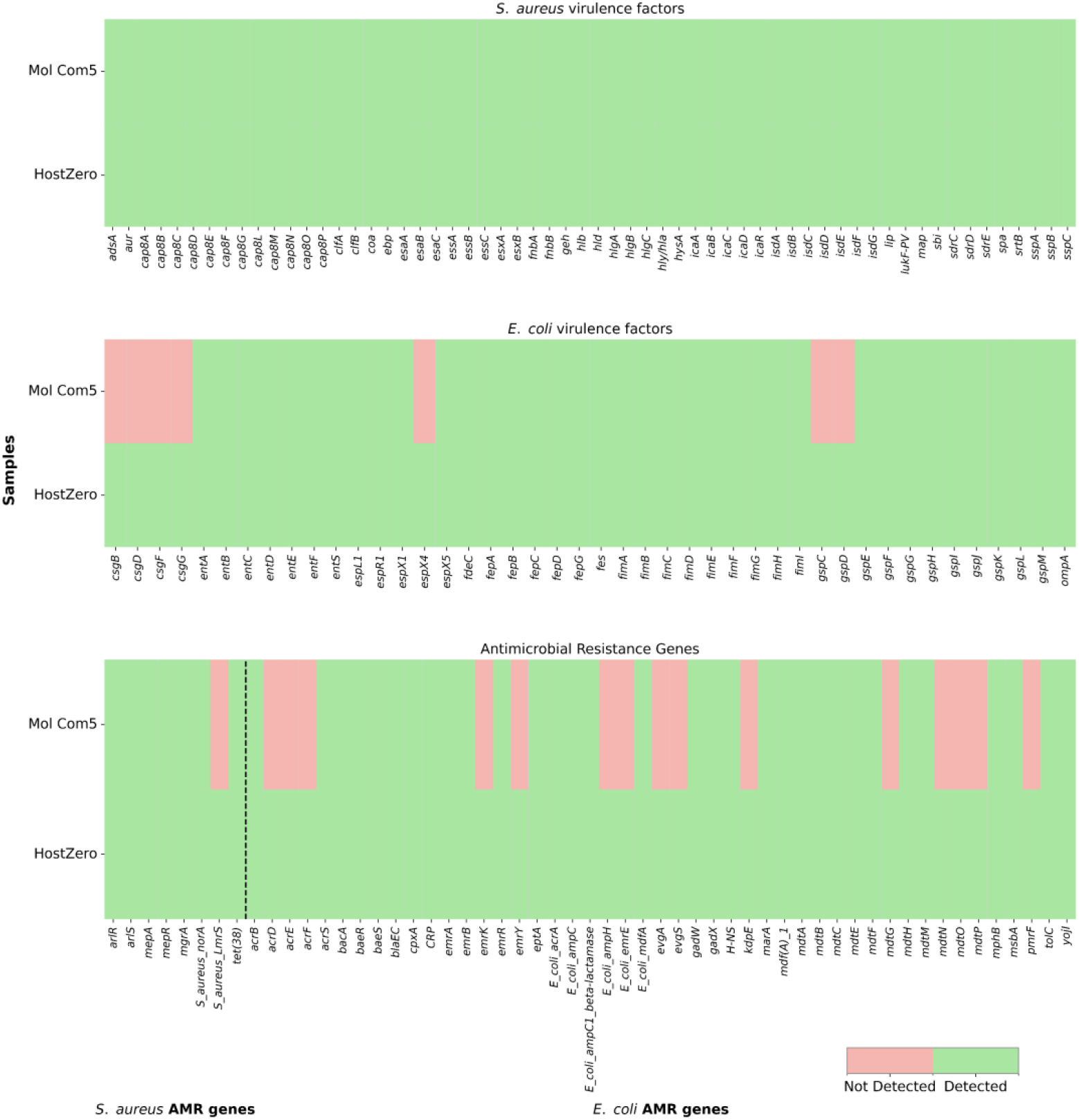
Heatmap of the AMR and VF genes detected through metagenomic sequencing of DNA extracted using the Mol Com5 and HostZero kits from milk samples. The panels in the figure represent (top) VF genes in the *Staphylococcus aureus* sample, (middle) VF genes in the *Escherichia coli* sample, and (bottom) AMR genes associated with both samples. Genes were considered detected (green) or not detected (red) based on alignment to VFDB, CARD, and NCBI resistance finder using ABricate.

## Discussion

Our previous study documented the successful identification of pathogens and detection of antibiotic resistance directly from mastitis milk samples, following optimization of a commercial kit and the use of long-read sequencing (Ahmadi et al., 2023). However, one of the main challenges in our previous work was enriching bacterial cells while minimizing inhibitory components such as proteins, lipids, and somatic cells present in the mastitis milk samples. To enrich bacterial cells, high-speed centrifugation is a commonly used method; however, prior studies (Stinson et al., 2021; Sun et al., 2019) have reported a significant proportion of bacteria trapped in the fat fraction after centrifugation of milk samples. We evaluated three pre-DNA extraction sample treatment methods across milk samples with varying bacterial loads (high, medium, and low CFU), aiming to recover bacterial cells trapped in the fat layer.

The supernatant of the milk samples after applying all three tested treatment methods contained some culturable bacterial cells. Using the standard centrifugation method (Method 1), overall bacterial recovery ranged from 82% to 95%, with greater losses in samples that had low initial bacterial concentrations. These results highlight the limitation of centrifugation in recovering all bacterial cells from milk samples, especially when the bacterial concentration is low. This observed bacterial loss in the fat layer is consistent with the findings of Sun et al. (2019), who reported a 7% loss in the fat layer under similar centrifugation conditions in milk samples spiked with 10^7^ CFU/ml of *S. aureus*. Similarly, Brewster & Paul (2016) reported that less than 7% of the bacteria in the pellet from raw milk were recovered following centrifugation, primarily due to bacterial entrapment within the fat layer. At higher bacterial loads, the binding capacity of the cream layer becomes saturated, resulting in improved recovery rates as a greater proportion of cells remain in the pellet (Brewster and Paul, 2016). A study conducted by Sinson et al. (2021) isolated both bacterial and human DNA from fat layers of the centrifuged human milk samples, highlighting that a significant amount of cells are trapped in the fat layer after centrifugation (Stinson et al., 2021). Although fat levels are typically lower in the infected milk samples (Bochniarz et al., 2023), the fat portion can vary considerably across samples, introducing variability in the efficiency of cell recovery by centrifugation.

In this study, to release the entrapped bacteria, the fat and whey fractions obtained after centrifugation were further treated with 0.1% Tween 20 and 2% citric acid. Tween 20 is a non-ionic surfactant commonly used to emulsify fats and oils in aqueous solutions (Frederick et al., 2013; Reichler et al., 2023). When combined with citric acid, an acidulant that clarifies the protein matrix (Seth & Bajwa, 2015), it improves microbial recovery from milk samples. The significant decrease in the CFU count in the supernatant after treating the fat and whey fractions with Tween 20 and citrate water (Figure 2A) suggests that chemical emulsification can release bacteria trapped in the fat layer. However, it introduces some uncertainty: the chemicals might have helped release the trapped bacteria into the pellet or could have affected bacterial viability. As a non-ionic surfactant, Tween 20 can disrupt bacterial membranes (Wu et al., 2013), potentially reducing cell culturability. Although we observed a lower CFU count in the fat layer after treatment, this does not guarantee the release of pathogens into the pellet or that a higher bacterial DNA yield will be achieved.

The use of gradient centrifugation can be advantageous for effectively separating cells in complex matrices, such as those found in milk. Previous studies (Fukushima et al., 2007; Meisel et al., 2011) have demonstrated the ability of Percoll gradients to effectively separate bacterial cells from milk and other complex food matrices. The near-complete absence of viable bacteria in the supernatant (Figure 2A) indicates efficient bacterial recovery. However, as with method 2, the potential impact of chemical treatment on cell viability complicates interpretation based solely on culturing.

We extracted DNA from the pellets obtained after treating aliquots of the same milk samples with all three methods and compared the bacterial and host Ct values. Although Method 1 led to the greatest loss of culturable bacterial cells (CFU count) in the supernatant, it consistently produced the lowest bacterial Ct and the highest host Ct, along with a balanced bacterial-to-host DNA ratio, as shown by qPCR analysis. This is important in applications like metagenomics or pathogen detection, where host DNA can dominate sequencing results and hide microbial signals. Methods 2 and 3 are more time-consuming, labor-intensive, and do not improve bacterial DNA recovery. Therefore, treating milk samples with Method 1 (centrifugation alone) is the preferred choice for microbial metagenomics and rapid diagnosis.

Host DNA depletion is a vital step in direct metagenomic sequencing of clinical samples because the high amount of host DNA can overshadow microbial DNA and interfere with accurate microbial profiling. In our previous study, we tested various DNA isolation kits specifically designed for microbial DNA extraction from food samples. However, pathogen and AMR identification was only successful with the Mol Com5 kit, which included additional micrococcal nuclease treatment to remove host DNA and enrich microbial DNA selectively (Ahmadi et al., 2023). In this study, we evaluated the performance of various commercial DNA kits with host depletion mechanisms in recovering bacterial DNA while minimizing host DNA contamination from bovine mastitis milk samples for downstream microbial metagenomics analysis. The Blood and Tissue kit, which lacks a selective host DNA depletion mechanism, served as a reference in this study. Although this kit produced higher DNA yield and better purity ratios, these results do not make it the best for metagenomics, as they reflect the total DNA from both host and pathogen, rather than specifically microbial DNA. Several studies (Ahmadi et al., 2023; Yap et al., 2020; Liu et al., 2023b; Bellankimath et al., 2024) have utilized DNA extraction kits without specific host depletion steps for milk and other clinical samples and reported that the majority of sequencing reads were of host origin. This high proportion of host DNA overshadows microbial DNA, which reduces the effectiveness of metagenomic sequencing in clinical diagnosis.

Three widely used kits for selective host DNA depletion and microbial DNA extraction were tested. The DNA yield was comparatively higher in samples with greater bacterial concentrations (CFU/ml) (Supplementary Table 2); however, the overall DNA yield remained low across all kits. The lower yield may result from the removal of host DNA, as the Blood and Tissue kit, which does not specifically deplete host DNA, consistently produced higher DNA yields in all samples. The fragment sizes and DNA integrity values of DNA extracted with the HostZero kit were notably greater than those from the Mol Com5 kit, indicating better preservation of high-molecular-weight DNA. The longer DNA fragments recovered by the HostZero kit are beneficial for metagenomic Nanopore sequencing, where longer DNA molecules support genome assembly and taxonomic classification (Quince et al., 2017; Maghini et al., 2021). No fragment size and DIN values were obtained from TapeStation for the DNA produced by the SPINeasy kit. This may be due to extremely low yield (Supplementary Table 2) or excessive fragmentation, potentially caused by suboptimal lysis or purification steps. Such degradation or loss of DNA significantly limits the usefulness of this kit for downstream metagenomic analysis, where both DNA quantity and integrity are crucial for accurate microbial profiling.

The qPCR-based assessment of bacterial and bovine DNA showed distinct differences in performance among the samples. The Blood and Tissue kit, as expected, yielded moderate bacterial Ct values but consistently low Ct values for the host target, indicating substantial presence of host DNA. Both the HostZero kit and Mol Com5 kit demonstrated consistent host DNA depletion across all tested samples, while preserving the bacterial DNA, which is supported by the findings of Marchukov et al.(2023). However, one should keep in mind that the complete depletion of host DNA is not achievable, and it heavily depends on the number of somatic cells and the change in sample composition due to infection (Marchukov et al., 2023). Samples with lower bacterial concentrations yielded very little DNA and had very high Ct values for both bacterial and host targets compared to samples with higher CFU. Studies have indicated that the number of somatic cells and bacterial count greatly affect both microbial and total DNA yield, with samples containing fewer somatic cells being more challenging for DNA extraction (Duarte and Porcellato, 2024). This indicates a need to develop a DNA extraction approach that effectively reduces host DNA while preserving bacterial cells to facilitate metagenomic diagnosis in samples with low bacterial content. The elevated Ct values for both bacterial and bovine targets in DNA extracted with the SPINeasy kit may result from inefficient extraction processes or excessively vigorous depletion steps, causing non-specific cell lysis and loss of DNA. This interpretation is reinforced by the notably low DNA yield from this method.

Samples with higher bacterial concentrations exhibited negative ΔCt values (bacterial Ct – host Ct) across all three host depletion kits, indicating efficient host depletion and a higher proportion of bacterial DNA (Supplementary Table 2). This further highlights the need to develop DNA extraction methods suitable for samples with low bacterial loads. Both the Mol Com5 kit and the HostZero kit demonstrated optimal and comparable performance in samples with higher CFU counts. However, in samples with lower CFU, the Mol Com5 kit showed a smaller ΔCt than the other tested kits, suggesting it has a better ability to enrich bacterial DNA in low-biomass samples. Despite this advantage, the Mol Com5 kit produced comparatively lower total DNA, which could limit its use in downstream sequencing applications. Additionally, the Mol Com5 kit effectively recovers DNA from gram-positive bacteria after host depletion. However, its performance with gram-negative bacteria was less optimal compared to the HostZero and Blood and Tissue kits (Supplementary Figure 3). Although only one gram-negative sample was tested in this study, our other experiments (unpublished results) consistently show reduced recovery of DNA from gram-negative pathogens using the Mol Com5 kit.

To overcome the limitation of low DNA yield, the ONT Rapid PCR Barcoding kit was used for library preparation, which is optimized for low input samples and requires less than 5 ng of starting DNA. Previous studies (Zhang et al., 2022; Simpson et al., 2023) have successfully used a PCR barcoding kit for low biomass samples. This kit involves PCR amplification of DNA, which produces amplicons of 2-5 kb (Oxford Nanopore Technologies, 2018). The comparable mean read lengths observed across all sequenced samples reflect the uniform amplicon sizes generated during the PCR, rather than inherent differences in initial DNA fragment sizes. However, a distinct impact of DNA integrity was observed on the number of reads and the quality of the assembly generated (Table 1 and Table 2). In this study, for DNA isolated using both HostZero and Mol Com5 kit, more than 75% of collected reads were assigned to the target pathogen, which is similar to the findings of (Ahmadi et al., 2023; Wright et al., 2023). The number of reads assigned to the target pathogen is slightly lower in Mol Com5.

Additionally, 5% reads were assigned to S. aureus in the sample infected with E. coli, which could lead to false-positive reports. Both samples sequenced in this study contained very high concentrations of bacteria (10^7^ CFU/ml). A limitation of this study is the small sample size and the focus on samples with high bacterial loads. The ability of these methods to detect pathogens and AMR genes in samples with lower bacterial concentrations using Nanopore sequencing remains to be tested. However, Grützke et al.,(2021) reported the identification of the pathogen using metagenomics shotgun sequencing from milk samples spiked with as low as 10^1^ CFU/ml of *Brucella abortus* was isolated with the HostZero kit. Although ONT offers adaptive sequencing, where only the DNA strand of interest is sequenced, we decided to disable this feature to gain a comprehensive understanding of the kit’s performance in host depletion and direct sequencing.

The Ct values for the bacterial target (17.6 and 17.2) and the bovine target (21.8 and 22) were similar in *S. aureus* infected samples, where DNA was extracted using both Mol Com5 and HostZero kits. The ΔCt (bacterial Ct – host Ct) was -4.2 for Mol Com5 and -4.8 for HostZero, a difference of only 0.6. Despite this small difference, HostZero produced about 10% more bacterial reads and 10% fewer bovine reads (Table 1) than Mol Com5. In contrast, for the *E. coli* sample, the bacterial Ct values were 18.8 (Mol Com5) and 14.1 (HostZero), and bovine Ct values were 25 (Mol Com5) and 19.3 (HostZero), resulting in ΔCt values of -6.2 (Mol Com5) and -5.2 (HostZero), a ΔCt difference of 1. This resulted in only 6% more target bacterial reads and 2% fewer bovine reads with the HostZero kit. These findings suggest that while ΔCt values from qPCR can give a rough estimate of host depletion and bacterial enrichment, they do not necessarily correlate proportionally with differences in sequencing read distributions.

Contiguity is essential for downstream genomic analysis, including taxonomic identification, AMR, and virulence factor detection in diagnostics, as well as structural variant detection and other analyses. Previous studies have shown that fragmentation during extraction adversely affects contiguity, affecting genome completeness and accuracy (Hillmann et al., 2018; Nicholls et al., 2019). Although a PCR barcoding kit was used in this study, which produces sequencing reads of similar length, a high number of reads and comprehensive genome assembly were achieved using the high-integrity DNA produced by the HostZero kit. The superior contiguity attained with HostZero supports its application in workflows requiring high-fidelity genome reconstruction.

These findings demonstrate that, with suitable DNA extraction and host depletion methods, direct metagenomic sequencing can achieve the same sensitivity and comprehensiveness of culture-based sequencing. Direct sequencing bypasses the need for bacterial cultivation, which is time and may fail to capture fastidious pathogens (Charalampous et al., 2019). Host DNA depletion is essential for high sensitivity. Consistent with our results, effective host depletion and microbial DNA enrichment with the HostZero kit have been documented before (Shi et al., 2022; Marchukov et al., 2023). Previous studies have shown that efficient host DNA depletion enhances the recovery of microbial DNA. This improves the ability to detect antimicrobial resistance genes and virulence factors, even when they are present at low levels in the clinical samples (Ahmadi et al., 2023; Bellankimath et al., 2025). Our results demonstrate that direct metagenomic sequencing, when optimized with effective host DNA depletion, can achieve the accuracy of culture-based genome sequencing for detecting AMR genes and virulence factors. The culture-independent approach not only reduces time to results but also minimizes biases associated with selective cultivation, making it a powerful tool for rapid pathogen characterization and antimicrobial resistance profiling in clinical settings.

## Conclusion

Sample preparation, which includes bacterial cell enrichment and DNA extraction, is crucial for culture-independent nanopore sequencing. This study shows that centrifugation alone is enough to enrich bacterial cells from milk samples without needing additional fat and whey fraction treatment with Tween 20 and citric acid. Additionally, effective host DNA depletion and microbial DNA enrichment are vital for diagnosing mastitis from milk samples. Among the tested methods with a selective host depletion mechanism, the HostZero kit proved to be the most effective in producing higher DNA with better integrity, which is beneficial for long-read sequencing and subsequent bioinformatics analysis. Ct values from qPCR provided insight into host depletion, which was reflected in sequencing; however, they may not directly correspond to the proportion of host and pathogen reads obtained from sequencing. This study confirms the use of a culture- and amplification-free metagenomic Nanopore sequencing approach to identify both gram-positive and gram-negative pathogens, as well as their antibiotic resistance profiles, in bovine mastitis milk samples, consistent with findings of Ahmadi et al. (2023). Furthermore, direct, culture-independent metagenomic sequencing provided AMR and virulence gene profiles comparable to those obtained from culture-based sequencing of isolates. Future studies will focus on sequencing a larger number of samples infected with different mastitis pathogens.

