## Supplementary Figure 1 for "Optimizing Sample Preparation for Direct Nanopore Sequencing to Enable Rapid Pathogen and Antimicrobial Resistance Profiling in Bovine Mastitis"

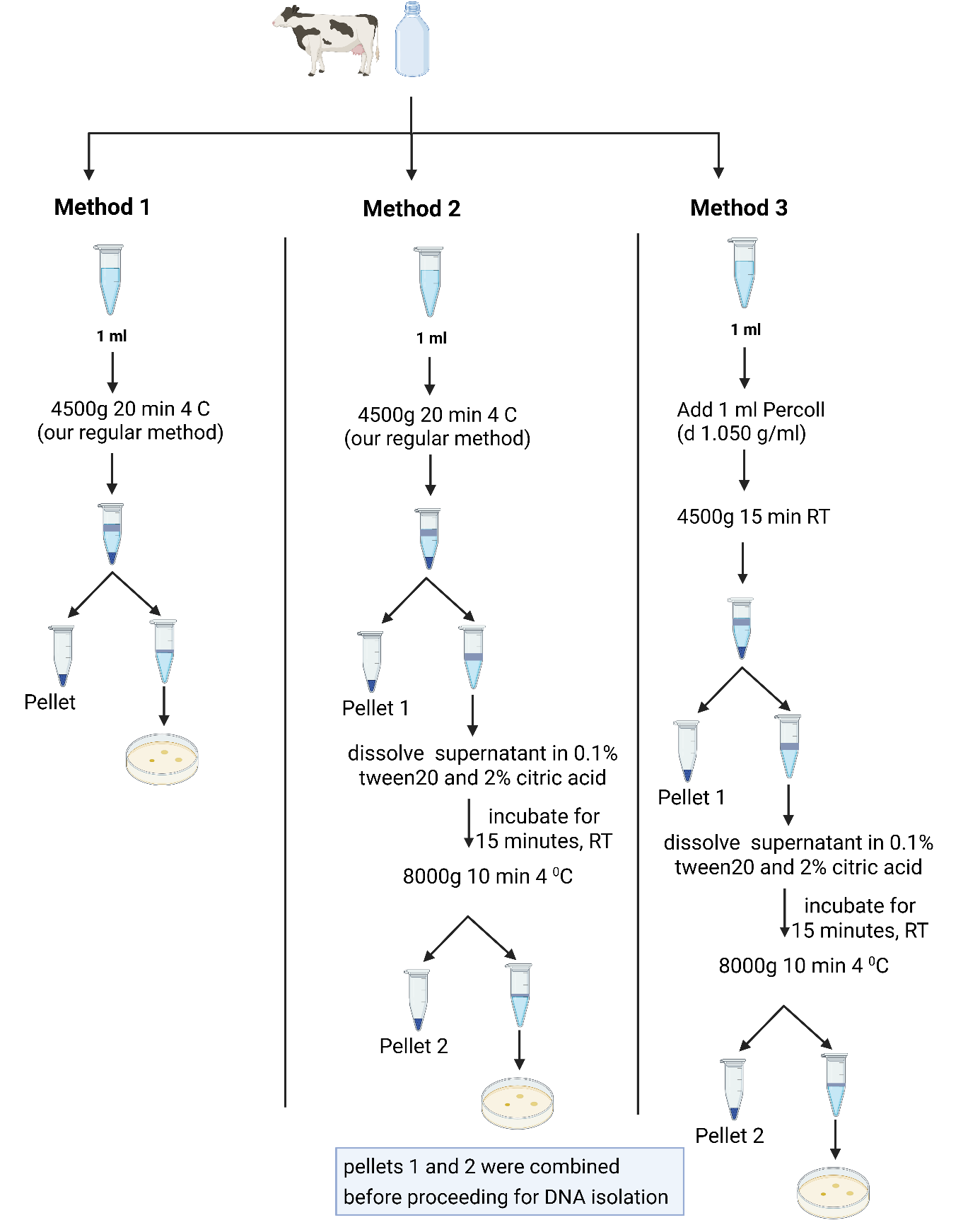

**Figure 1. Experimental design of pre-DNA extraction sample treatment methods.** Three 1 ml aliquots of the same milk sample were processed using Method 1, 2 and 3. 100 ul of the final supernatant that was discarded was cultured in BHI agar to count the CFU loss. All the pellets were subjected to DNA extraction (in the case of methods 2 and 3, the pellets 1 and 2 were merged) using Mol Com5_cent-nuc_ protocol.
