## Supplementary Figure 2 for "Optimizing Sample Preparation for Direct Nanopore Sequencing to Enable Rapid Pathogen and Antimicrobial Resistance Profiling in Bovine Mastitis"

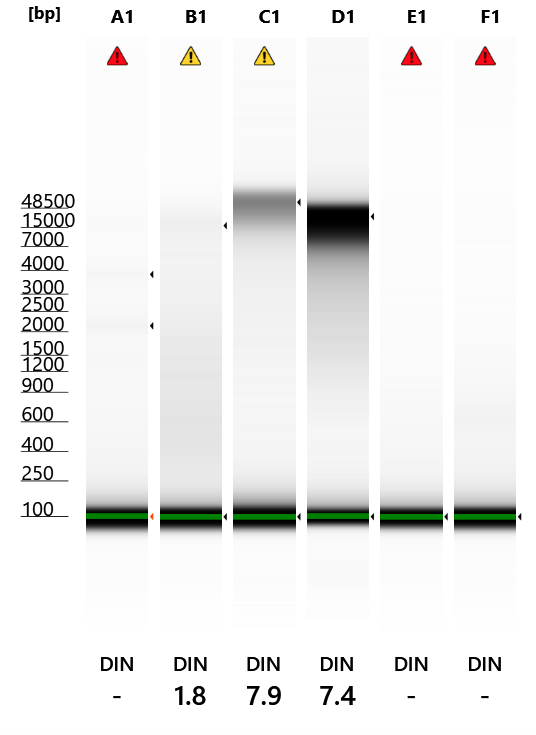

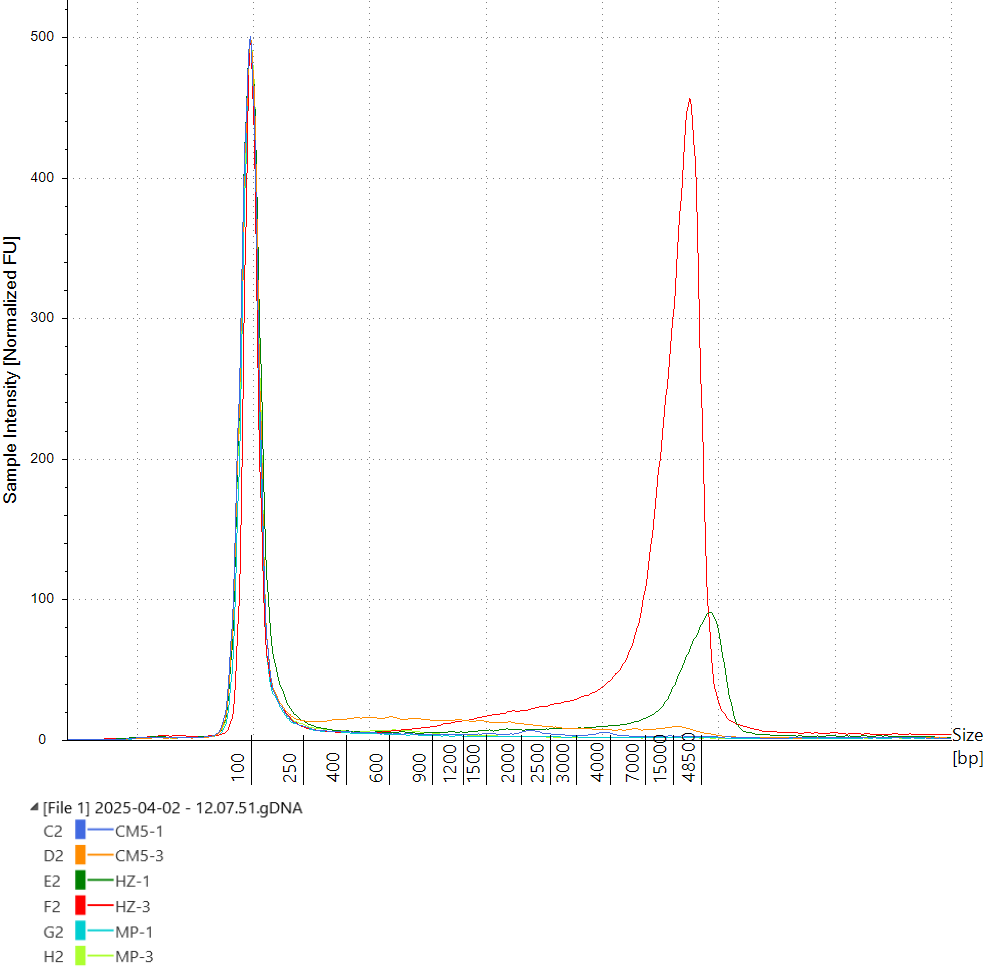

**B**

**A**

**Figure 1.** Assessment of DNA quality and fragment size distribution using Agilent TapeStation system, genomic screen tape. **A.** Sample integrity plot; CM5 : Mol Com5 kit, HZ: Host Zero kit, MP: SPINeasy kit; 1: S. aureus sample, 3: E. coli sample. **B.** Corresponding gel-like image from Agilent TapeStation. Lanes A1 and B1: Mol Com5 S. aureus and E. coli, C1 and D1: HostZero S. aureus and E. coli, E1 and F1: SPINeasy S. aureus and E. coli. Black triangular dots in the lanes represent bands with the highest intensity. HostZero kit has the largest fragment sizes as well as a better DNA integrity (DIN) values. There was no DNA detected in lanes E1 and F1 (DNA extracted with SPINeasy kit).
