## Supplementary Figure 3 for "Optimizing Sample Preparation for Direct Nanopore Sequencing to Enable Rapid Pathogen and Antimicrobial Resistance Profiling in Bovine Mastitis"

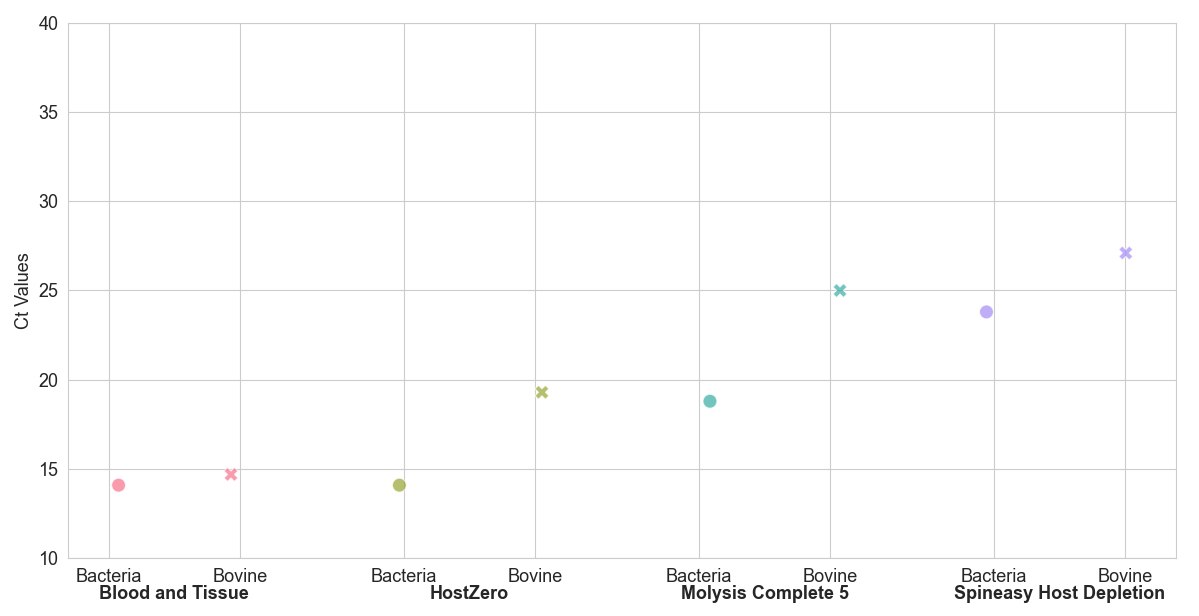

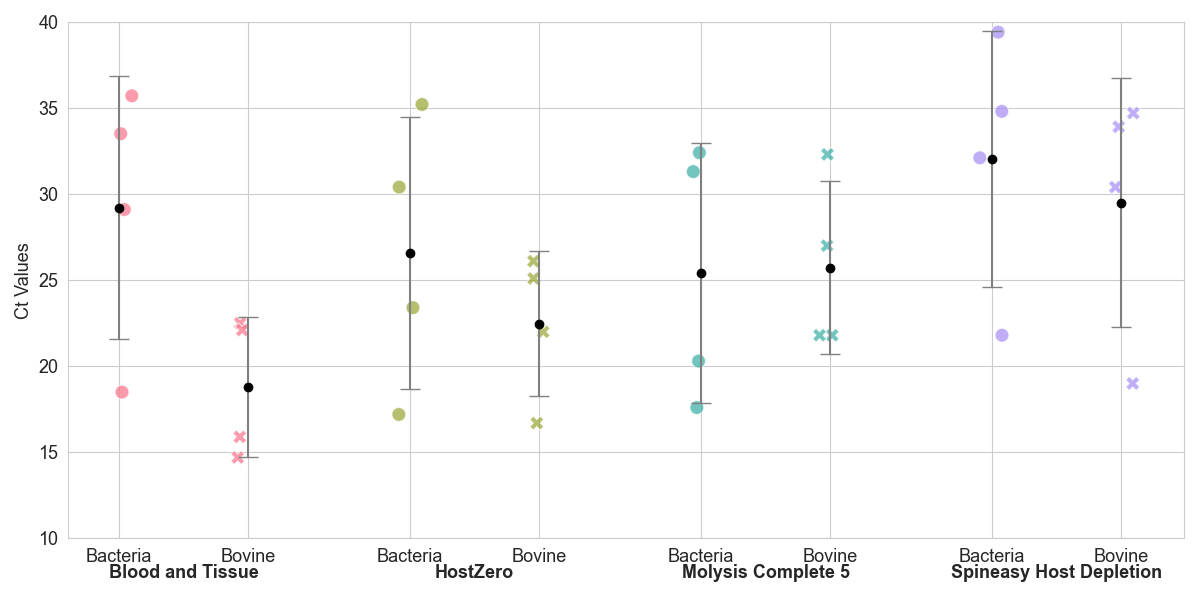

dots**Figure 1.** Ct values plot for bacterial target and bovine target in the DNA extracted using different kits from milk samples. **A.** Samples infected with gram-positive pathogens: dot and cross represent the Ct values for each sample. **B.**  Sample infected with gram negative E. coli.

**B**

**A**
